# A virion protein of an archaeal virus inhibits a Type V BREX defense system in *Haloarcula hispanica*

**DOI:** 10.64898/2026.04.28.721311

**Authors:** Julia Gordeeva, Richard D. Morgan, Ananya Jain, Alexey Fomenkov, Yvette Luyten, Tamas Vincze, Richard J Roberts, Alfred A. Antson, Konstantin Severinov

## Abstract

Bacteriophage Exclusion (BREX) systems are a diverse family of bacterial and archaeal defense mechanisms defined by two conserved genes encoding a putative alkaline phosphatase BrxZ (PglZ) and an ATPase BrxC (PglY). Here, we characterize a Type V BREX system from the archaeon *Haloarcula hispanica*. Similar to bacterial Type I BREX systems, host–virus discrimination relies on methylation of specific non-palindromic DNA motifs. Notably, the *H. hispanica* BREX locus encodes two methyltransferases, BrxX1 and BrxX2, which independently target distinct motifs (GTAYCCG and GACCCC). The system protects *H. hispanica* against three of seven tested archaeal viruses. However, two related viruses, SH1 and HHIV-2, despite encoding multiple BREX target sites, escape BREX-mediated defense. We demonstrate that a large virion protein, VP1, encoded by these viruses inhibits the BREX system, as mutant viruses carrying partial deletions of VP1 lose resistance to the host defense. Together, these findings provide the first characterization of a Type V BREX system and demonstrate that archaeal viruses can counteract BREX through virion-associated anti-defense proteins.

## INTRODUCTION

The continuous evolutionary conflict between prokaryotes and their viruses has driven the emergence of prokaryotic immune systems and, in turn, viral anti-immune mechanisms. One defense family, BREX (Bacteriophage Exclusion), is found in both bacterial and archaeal genomes and comprises systems centred around two core genes encoding a putative alkaline phosphatase BrxZ (PglZ) and a large protein BrxC (PglY) with an ATP-binding domain (1,2). The family is classified into six types based on the composition of additional genes within the BREX loci (1). The most common and best-studied are Type I BREX systems (1,3–6).

The recognition strategy of Type I BREX resembles that of restriction–modification systems: host DNA is protected by methylation at a specific motif (BREX site), whereas invading phage DNA that lacks this modification is recognized as foreign. The methyltransferase BrxX alone lacks detectable methylation activity but determines the BREX site recognition and potentially serves as a sensor of phage infection (7). BREX site methylation is mediated by a multiprotein complex consisting of BrxX, BrxC, BrxZ, and an additional small protein BrxB (3,7). Unlike the classical restriction-modification systems, BREX loci do not encode typical restriction endonucleases. Though BrxZ was recently shown to be a metal-dependent nuclease, *in vitro* it cleaved both methylated and non-methylated dsDNA (8). Other components of Type I BREX, including a DNA-binding protein BrxA (9) and an AAA+ ATPase BrxL (10). Type II BREX systems were originally discovered as the phage growth limitation (Pgl) system in *Streptomyces coelicolor* (11,12). The system consists of four components including conserved proteins PglZ and PglY, a kinase PglW and a SAM-dependent DNA methyltransferase PglX (11). The proposed mechanism of action differs from that of classical restriction–modification systems and involves epigenetic marking of phage DNA during the first round of infection, thereby flagging it for restriction in subsequent infection cycles (12,13). The Pgl-mediated defense is thought to rely on a toxin–antitoxin– like mechanism in which the toxic activity of PglX is controlled by PglZ; the phage infection triggers the formation of modifying or restricting complexes, with the latter inhibiting phage infection (11).

Other BREX types (III–VI) have not been functionally characterized. In this work, we explore a Type V BREX system found exclusively in archaeal genomes (1). We demonstrate that the recognition strategy of a Type V BREX from archaeon *Haloarcula hispanica* resembles that of the bacterial Type I systems: the host genome is protected by methylation of specific BREX sites while invader’s DNA is targeted by BREX. We found that some archaeal viruses with multiple BREX sites in their genomes escape the BREX defense. We demonstrate that these viruses encode a large virion protein with an anti-BREX activity.

## MATERIALS AND METHODS

### Strains and growth conditions

Strains and viruses used in this study are listed in Supplementary Table S1. *Haloarcula hispanica* ATCC 33960 *ΔpyrF* and its derivatives were generally cultivated in modified growth medium (MGM) supplemented with 50 μg/ml uracil at 37°C. Artificial, 30% saltwater (SW) containing per litre 240g NaCl, 30g MgCl_2_×6H_2_O, 35g MgSO_4_×7H_2_O, 7g KCl and 5 ml of 1M CaCl_2_×2H_2_0 was prepared as described in the Halohandbook (14). One litre of MGM contained 5g of peptone (Oxoid), 1g of Bacto yeast extract (Oxoid), and 23% SW for liquid media, 18% SW and 4g of agar (Oxoid) for top-layer or 20% of SW and 14g of agar for solid media, pH 7-7.5. For transformation selection, *H. hispanica* cells were plated on yeast extract-subtracted AS-168 (AS-168YE-) (15). Per liter, 200g NaCl, 20g MgSO_4_×7H_2_O, 2g KCl, 3g trisodium citrate, 1g sodium glutamate, 50 mg FeSO_4_×7H_2_O, 0.36 mg MnCl_2_×4H_2_O, 5g Bacto casamino acids (BD), pH 7-7.5. *Escherichia coli* strains were grown in Luria-Bertani (LB) medium supplemented with ampicillin added to a final concentration of 100 mg/ml and used for cloning.

### Plasmid constructions

For deletion of the whole BREX^HAR^ system (NC_015943.1, positions 401 474 - 426 563), a 600 bp upstream region of *brxA* and a downstream sequence of *brxHII* were amplified with primers US_ΔBREX_F/R and DS_ΔBREX_F/R, respectively (Supplementary Table S2). Two PCR fragments were merged by overlap extension PCR, treated with KpnI and BamHI restriction enzymes, ligated with the suicide plasmid pHAR and transformed into *E. coli* DH5α. For the construction of a partial deletion of BREX^HAR^ (deletion of the fragment *brxX1-brxZ-brxX2-brxHII*), the same procedure was repeated with primers US_ΔX1-HII_F/R for amplification of the upstream region and DS_ΔX1-HII_F/R for amplification of the downstream region. Deletions of individual genes, pHAR-ΔN, where N = *brxA, B, C1, C2, X1, X2, Z, HII, nucS*, were created with primers US_ΔN_F/R and DS_ΔN_F/R.

Plasmids for an anti-BREX gene search was created by amplification of viral DNA fragments of SH1 and HHIV2 viruses using Phusion Plus polymerase (Thermo Scientific) with primers HHIV2_Pn_InF_F/R or SH_P2_InF_F/R, respectively, each containing 15 bp overlaps to pWL502, where n = 1–7. To introduce stop codons, two DNA fragments were generated using primer pairs SH1/HHIV2_P2_InF_F and SH1/HHIV2_P2_XXStopXX_R, as well as SH1/HHIV2_P2_InF_R and SH1/HHIV2_P2_XXStopXX_F. To clone pWL502-HHIV2_P2_VP1 plasmid *HHIV2_gp5* gene were amplified with HHIV2_P2_InF_F and HHIV2_P2_VP1_R primers. To clone pWL502-SH1_P2_VP1/gp13-16 plasmids, fragments were generated using SH1_P2_gp13_F and SH1_P2_gp13_R/SH1_P2_InF_R primers, respectively. The archaeal vector pWL502 was linearized using primers pWL502_lin_F and pWL502_lin_R. Ligation of DNA fragments and the vector was performed using the In-Fusion cloning method (Takara), and the resulting constructs were transformed into *E. coli* Stellar competent cells (Takara). Subsequently, *H. hispanica* DF60 Δcas1-8 BREX+/ΔBREX cells were transformed with the plasmids and plated on AS169YE-medium.

CRISPR targeting plasmid was generated by the In-Fusion reaction of the linearized pWL502 and two fragments of the CRISPR array created by annealing primers CRISPR_array_F1/R1 and CRISPR_array_F2/R2. The resulting plasmid, containing a *phaR* promoter and two CRISPR repeats, was linearized with pWL502_crRNA_lin_F2/R2 and merged with 400 bp upstream and downstream regions of *HHIV2_gp5* amplified with primers US_HHIV2_gp5_F/R and DS_HHIV2_gp5_F/R. The spacer_N sequence, where N= -1, 0, 1, 2, 3 and 4, was created by annealing primers spacer_N_F/R and inserted into the linearized plasmid with the CRISPR array and the upstream and downstream sequences of *HHIV2_gp5* amplified with pWL502_crRNA_lin_F/R. All constructions were transformed into *E. coli* Stellar competent cells and subsequently into *H. hispanica* DF60 ΔBREX with the active CRISPR-Cas system.

### Gene knockouts in the BREX^HAR^ locus

Gene knockouts in uracil auxotrophic *H. hispanica* DF60 strain (*ΔpyrF)* with deleted *cas* genes were created by a pop-in-pop-out method (15). The suicide plasmids containing upstream and downstream regions flanking the desired gene were transformed in *H. hispanica* by a protocol described in the Halohandbook (14). Transformants were plated on AS168YE-to positively select the single-crossover recombinants (pop-in). The double-crossovers were selected by cultivation of cells in AS168 medium supplemented with uracil (50 μg/ml) and 5’-FOA (150 μg/ml) to excise a plasmid. The mutants were screened by PCR analysis and sequenced.

### Efficiency of plaquing (EOP) assay

Late exponential phase cultures of *H. hispanica* DF60 (150 μl) were mixed with 10 μl of viral lysates or their serial (10^−1^ -10^−8^) dilutions and 2.5 ml of 18% MGM soft agar supplemented with uracil (for pWL502-deficient cells) and poured on the surface of precast 20% MGM plates. After 3-5 days of incubation at 30 - 37°C depending on virus preference, the efficiency of plaquing was determined as a ratio of virus titers on BREX+ to BREX-lawns.

### Growth curves of infected cultures

Late exponential phase cultures of *H. hispanica* DF60 were diluted 1:50 in 23% MGM supplemented with uracil and grown at 37 °C to OD_600_ = 0.6-0.8. At t = 0 ΔBREX and BREX+ cultures were infected with HHPV3 virus to reach MOI = 1. At t = 3 hours post-infection unabsorbed viruses were washed away by centrifugation at 10.000 × g for 15 min and resuspension in the original volume of fresh MGM with uracil. The dynamics of infected cultures growth was monitored by measuring OD_600_ every 30 min. At various times post-infection (0, 3, 4, 5, 6, 7, 8, 9 and 10 hours) aliquots from infected cultures were taken to determine virus titer (PFU) and the number of living cells (CFU).

### SMRT Motif and Modification analysis of *H. hispanica* genomes

Archaeal genomic DNAs were purified using Genomic DNA Purification kit (Thermo Scientific) and the DNA sample was sheared to an average size of ∼10 kb using the G-tube protocol (Covaris, MA). DNA libraries were prepared using a SMRTbell express template prep kit 2.0 (100-938-900, Pacific Bioscience, CA) and ligated with hairpin barcoded adapters lbc. Partially formed SMRTbell templates and unligated SMRTbell adapters were removed by digestion with a exonuclease III and exonuclease VII (NEB, MA). The qualification and quantification of the SMRTbell libraries were made on a Qubit fluorimeter (Invitrogen, OR) and a 2100 Bioanalyzer (Agilent Technologies, CA). SMRT sequencing was performed using an SQ1 (Pacific Biosciences, CA) based on the multiplex protocol for 10 kb SMRTbell library inserts. Sequencing reads were collected and de novo assembled using the Microbial Assembly version 10.1.0.1119588 program with default quality and read length parameters. In addition to genome assembly (16), the SMRT Analysis pipeline from Pacific Biosciences (http://www.pacbiodevnet.com/SMRT-Analysis/Software/SMRT-Pipe) enables the determination of the epigenetic status of sequenced DNA by identifying the m6A and m4C modified motifs (17–19).

### CRISPR targeting of HHIV2 virus

Late exponential phase cultures of *H. hispanica* DF60 ΔBREX with the active CRISPR-Cas system and pWL502 plasmid with a spacer targeting *HHIV2_gp5* gene (150 μl) were mixed with 100 μl of HHIV2 virus (10^10^ PFU/ml) and 2.5 ml of 18% MGM soft agar and poured on the surface of precast 20% MGM plates. After 5-7 days of incubation at 30°C, escaper plaques were picked and produced on DF60 Δ*cas1-8* ΔBREX lawn. DNA of escapers was purified by phenol-chloroform extraction and amplified by ΔVP1_scrn_F/R primers flanking *HHIV2_gp5* gene. DNA fragments were sequenced to identify mutations.

### HHIV2 virus production and purification

Virus was precipitated from collected plate lysates of semi-confluent top-layer agar plates with 10% (w/v) polyethylene glycol 6000 MW (PEG), pelleted by centrifugation at 18,000 rpm for 60 min at 4 °C (Sorvall SS34), and resuspended in HHIV2 buffer (20 mM Tris-HCl pH 7.2, 20 mM MgCl_2_, 10 mM CaCl_2_, and 0.5 M NaCl)(20). The solution was applied to the top of a caesium chloride step gradient in an Ultra-Clear tube (Beckman), comprising solutions of densities 1.25, 1.35 and 1.41 g/ml in HHIV2 buffer. Gradients were centrifuged at 28 000 rpm in a Himac P40ST rotor for 17 hours. The light-scattering virus zone was collected and pelleted with 38% sucrose cushion (HHIV2 buffer) at 32 000 rpm for 3 hours (Himac P40ST rotor).

### Protein analysis

HHIV2 virion composition was analysed in a 4–12% gradient Bis-Tris SDS gel (NuPAGE, Invitrogen) stained with Coomassie brilliant blue. Virus proteins were identified by MALDI-MS/MS analysis of the tryptic peptides in the Centre of Excellence in Mass Spectrometry, Technology Facility, University of York.

## RESULTS

### Type V BREX systems recognition strategy is similar to that of Type I systems

Type V BREX systems were found only in archaea belonging to the *Halobacteria* class (1). The Type V BREX system from *Haloarcula hispanica* ATCC 33960 (BREX^HAR^) shares several genes with BREX^Ec^, a well-studied Type I system from *E. coli* (3) (Fig. 1a). Both systems encode an HTH-domain protein BrxA; a protein of unknown function BrxB, and a nuclease BrxZ. While *E. coli* BREX encodes a single BrxC (PglY) ATPase and BrxX (PglX) DNA-methyltransferase, the *H. hispanica* system encodes two versions of each protein. BrxC1 and BrxC2 share 13% sequence identity with each other whereas BrxX1 and BrxX2 sequences are 53% identical. Additionally, both systems encode AAA+ domain proteins, BrxL in *E. coli* and BrxHII in *H. hispanica*, with no detectable sequence identity. The archaeal protein is predicted to be a helicase.

**Figure 1.**
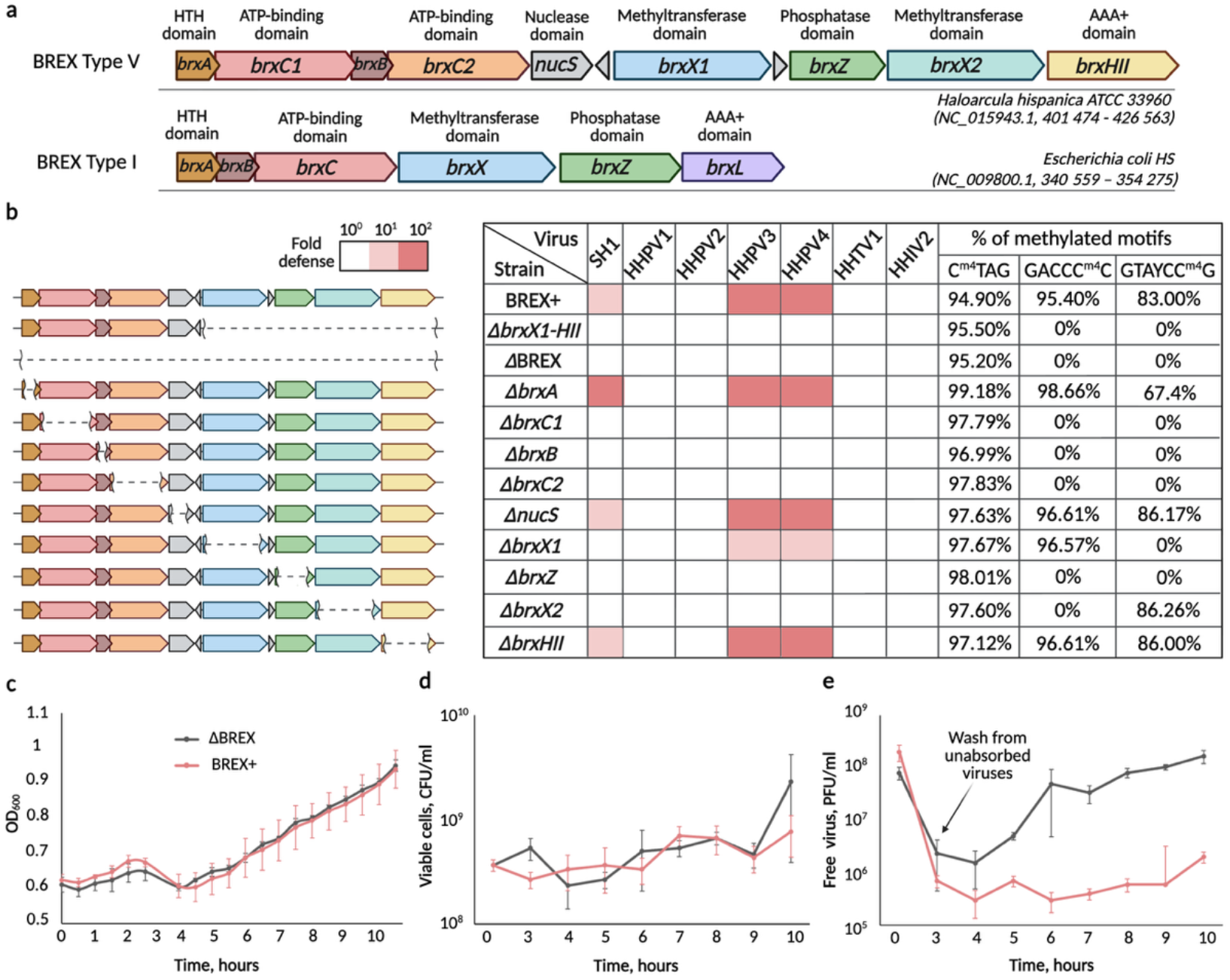
Type V BREX system from *Haloarcula hispanica* limits infection by diverse viruses. **a**. Comparison of the Type I BREX system from *E. coli* HS and Type V BREX system from *H. hispanica*. **b**. Defense profiles of *H. hispanica* wild-type and indicated BREX mutant strains against a collection of *H. hispanica* viruses and statistics of methylated DNA sites in the genomes of tested strains. Fold defense was measured using serial dilution plaque assays, comparing viral titers on strains carrying indicated deletions in the BREX^HAR^ locus to the titer on a control ΔBREX strain lacking the entire locus. Data represent averages of three replicates. **c**. Growth curves of BREX+ and ΔBREX cultures infected with the HHPV3 virus at a multiplicity of infection (MOI) of 1. Infection was initiated at time = 0. Each curve represents the mean of three independent experiments. **d**. Colony-forming units (CFU) in BREX+ and ΔBREX cultures from panel **c**. **e**. Plaque-forming units (PFU) in BREX+ and ΔBREX cultures from panel **c**.

To investigate BREX^HAR^-mediated defense, we used an uracil auxotrophic (*pyrF*-deleted) *H. hispanica* strain DF60 with inactivated CRISPR-Cas system (21). We will refer to this strain as BREX+. As controls, we created isogenic strains with complete (ΔBREX) or partial (Δ*brxX1-HII*) deletions of the BREX^HAR^ locus. We assumed that similar to BREX^Ec^, BREX^HAR^ functions by modifying and protecting the host genome and eliminating non-modified invader’s DNA (3). Therefore, *H. hispanica* viruses produced upon infection of a strain with deficient BREX should not contain BREX-mediated modifications and should be subject to exclusion by the BREX+ host. Seven *H. hispanica* viruses from laboratory collection were propagated on the ΔBREX strain and considered as lacking BREX modifications. The efficiency of plaque formation (EOP) of ‘unmodified’ viruses on BREX+ cells lawns was compared to that on ΔBREX, and Δ*brxX1-HII* control lawns. As seen from Fig. 1b, the BREX+ strain was ∼100-fold more resistant to HHPV3 and HHPV4 viruses, and ∼10-fold more resistant to the SH1 virus than the control strains.

The growth rates of ΔBREX and BREX+ cultures infected with the HHPV3 virus were next monitored. Liquid *H. hispanica* cultures were infected with HHPV3 and unabsorbed viruses were washed away three hours post-infection. The HHPV3 virus does not lyse infected cells (22) and, therefore, no decline in colony forming units (CFU) or in optical density (OD_600_) of infected cultures was observed, as expected (Fig. 1c, d). In the ΔBREX culture, the HHPV3 progeny release was detected starting ∼6 hours post-infection (Fig. 1e). In contrast, no increase in viral titer was observed in the infected BREX+ culture. We conclude that similar to the Type I BREX^Ec^, Type V BREX^HAR^ prevents the production of progeny by sensitive viruses.

To map the essential components of BREX^HAR^, we engineered a series of *H. hispanica* strains with deletions of individual BREX genes. Deletions of *brxA* and *brxHII* genes had no effect on BREX^HAR^ defense against either HHPV3 or HHPV4 (Fig. 1b). Curiously, deletion of *brxA* showed ∼10-fold increased protection against SH1 compared to that observed in cells with the wild-type BREX locus. Deletions of *brxB, brxC1, brxC2, brxX1, brxX2*, and *brxZ* abolished protection against HHPV3, HHPV4, and SH1. In addition to canonical BREX genes, a gene encoding a predicted endonuclease NucS (nuclease for ssDNA) is inserted between the *H. hispanica brxC2* and *brxX1* genes (Fig. 1a). A similar insertion of a *brxU* endonuclease gene is present in the Type I BREX system from *Escherichia fergusonii* (4). However, cultures of *H*.*hispanica* with deletion of *nucS* remained fully protected against viral infection.

Genomic DNA of *H. hispanica* BREX+ and mutant strains was sequenced using Single Molecule Real-Time sequencing (SMRT). In all cases, the CTAG sites contained N^4^-methylcytosines on both strands (Fig. 1b), presumably a result of modification by a CTAG-methyltransferase, the product of the *zim* gene (23). CTAG motives methylated by different DNA methyltransferases M.HhiI are common among most halophilic archaea (24–26). The CTAG sequences are strongly underrepresented in *H. hispanica* viruses (27), suggesting that there may be a host defense mechanism that relies on CTAG site methylation for self/non-self-differentiation. N^4^-methylation of cytosine at the fifth position of GACC<u>C</u>C sites and cytosine at the sixth position of GTAYC<u>C</u>G (Y = C or T) motifs was found in DNA from BREX+ but not from ΔBREX cells. These modifications were also detected in DNA of *brxA, nucS*, and *brxHII* mutants. Deletion of *brxX1* abolished methylation of GTAYCC^m4^G sites, while deletion of *brxX2* resulted in the absence of the GACCC^m4^C modification. Disruption of either *brxC1* or *brxC2* abolished methylation of both sites, suggesting that *H. hispanica* BREX modification complexes require both ATPases to function.

### Identification of an anti-BREX gene

The *H. hispanica* viruses used in this study were analysed for the presence of GACCCC and GTAYCCG motifs modified by BREX^HAR^ (Fig. 2a). Although the number of BREX*-*recognized sites in closely related viruses SH1 and HHIV2 is higher than in HHPV3 and HHPV4, the level of BREX protection against SH1 is weaker than that against HHPV3 and HHPV4. Further, no protection against HHIV2 is observed. We hypothesized that HHIV2 may encode a protein that deactivates the BREX defense. The entire genome of HHIV2 was subcloned, as seven independent fragments, into the pWL502 plasmid vector, producing plasmids HHIV2_Pn, where n = 1 to 7 (Fig. 2b). Because of toxicity, we were unable to clone *gp18*, a gene encoding a structural protein VP13 with a transmembrane domain (20), and it was thus omitted from plasmids HHIV2_P2 and HHIV2_P3. HHIV2_Pn plasmids were individually transformed into either ΔBREX or BREX+ *H. hispanica* cells and the level of BREX^HAR^ defense of plasmid-bearing cells against HHPV3 and HHPV4 was determined (Fig. 2c).

**Figure 2.**
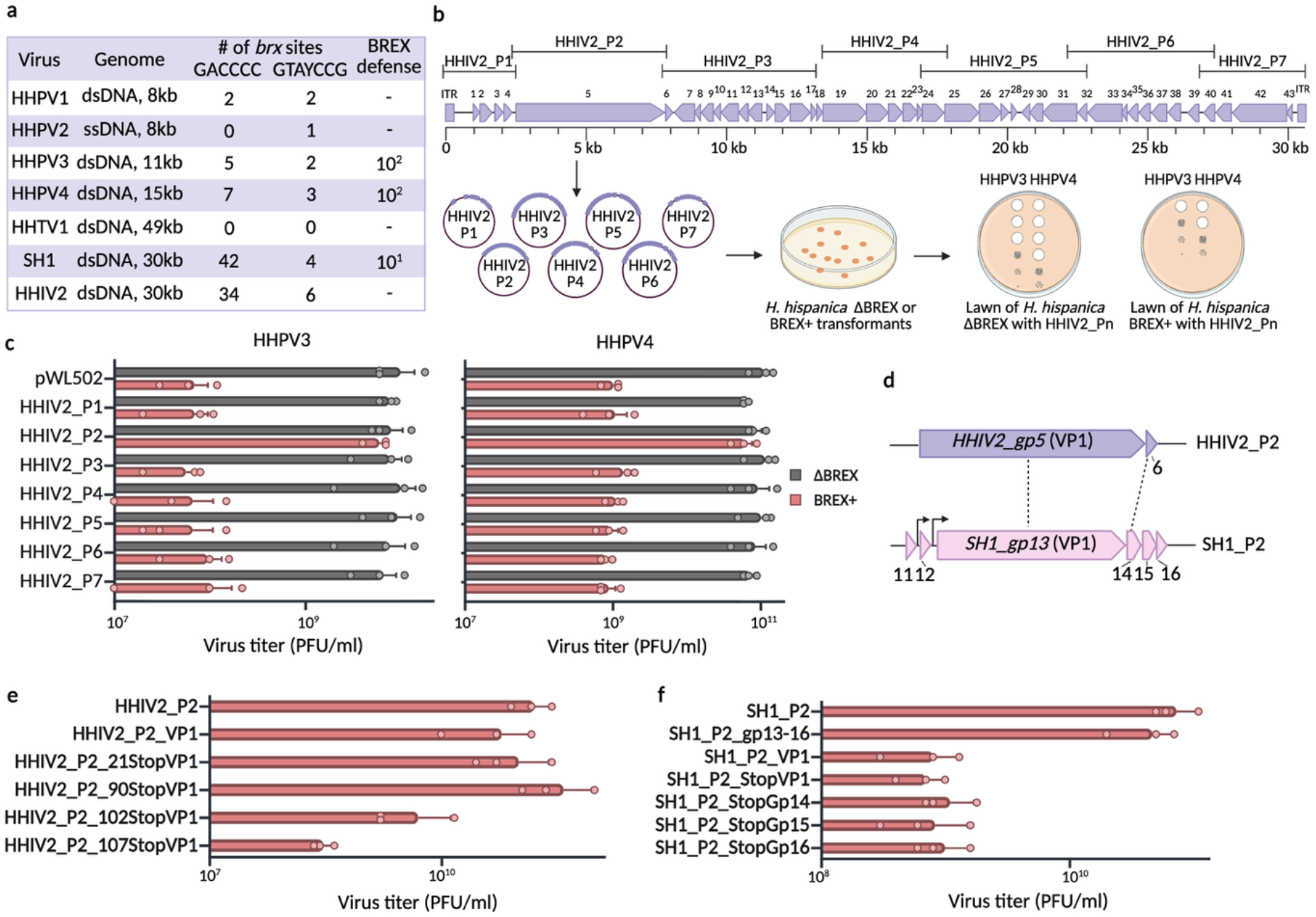
Anti-BREX proteins encoded in HHIV2 and SH1 viruses. **a**. The number of BREX sites in haloarchaeal viruses, viral genome sizes, and the levels of BREX defense against these viruses. **b**. Schematic representation of anti-BREX proteins identification experiment by cloning the HHIV2 genome into seven separate plasmids, followed by transformation into ΔBREX or BREX+ *H. hispanica* strains. **c**. Effect of different HHIV2 genome fragments on BREX defense against HHPV3 and HHPV4. **d**. Genes and proteins encoded by the HHIV2_P2 and SH1_P2 plasmids. **e**. Effect of different HHIV2_P2 plasmid variants on BREX defense activity against HHPV4. **f**. Effect of different SH1_P2 plasmid variants on BREX defense activity against HHPV4.

The presence of the HHIV2_P2 plasmid abolished the BREX defense against both HHPV3 and HHPV4 (Fig. 2c). This plasmid contains *gp5*, a gene that encodes the largest HHIV2 protein VP1 (20), and a fragment of a short downstream *gp6* gene. The VP1 protein is part of the HHIV2 virion (20) but it has not been identified in the virion structure obtained by cryo-EM (28,29), suggesting that it may be an internal protein closely associated with tightly packed viral DNA. The *gp13* gene of SH1 is a homolog of HHIV2 *gp5* (20) (Fig. 2d). Plasmid SH1_P2 that contained the SH1 *gp13* gene and several short upstream and downstream genes also inhibited the BREX defense (Fig. 2f).

To determine whether the VP1 protein is necessary for BREX inhibition, the HHIV2_P2 plasmid derivatives with stop codons introduced in the beginning of HHIV2 *gp5* and SH1 *gp13* were prepared (HHIV2_P2_21StopVP1 and SH1_P2_StopVP1, respectively). Stop codons introduced in *SH1_gp13* but not in *HHIV2_gp5* abolished the inhibition of BREX defense (Fig. 2e, f). The N-terminal sequence was experimentally confirmed only for SH1 VP1 (20,30). We hypothesised that the start codon of HHIV2 VP1 is misannotated and that translation initiates not from a methionine codon at the annotated position 1 but from the next methionine codon at the annotated position 103 (Fig. 3b). Consistent with this idea, a stop codon introduced instead of a threonine codon at position 107 of *HHIV2_gp5* abolished BREX defence inhibition, while a stop codon introduced instead of a leucine codon at position 90 had no such effect (Fig. 2e, 3b). A stop codon introduced instead of a glycine codon at position 102 resulted in partial inhibition of BREX defense, potentially because of the alteration of a Shine–Dalgarno sequence in front of Met103, which, based on our data, acts as a start codon of HHIV2 VP1 and should be considered as Met1.

**Figure 3.**
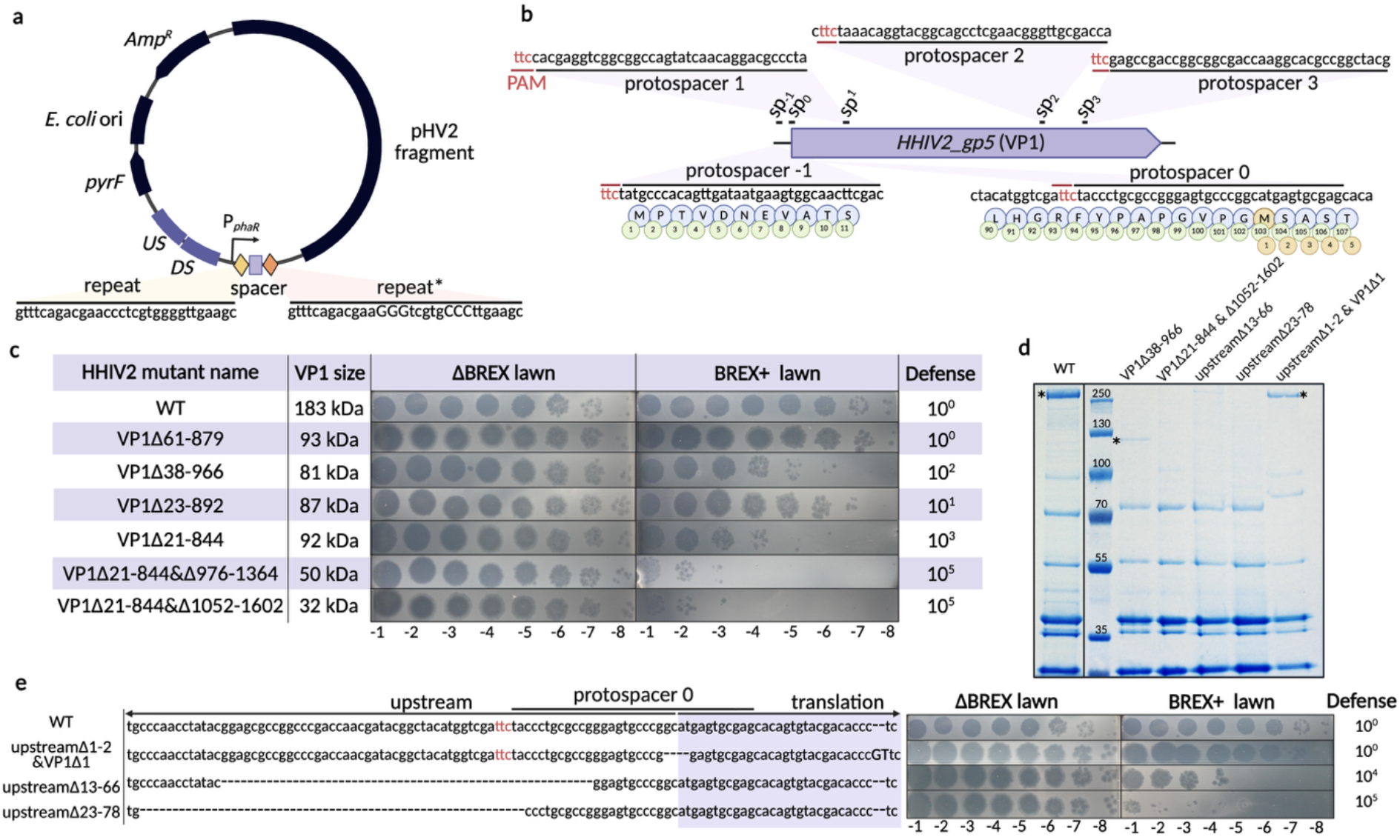
Effects of mutations in VP1 on HHIV2 ability to overcome the BREX defence. **a**. Schematic representation of a plasmid carrying a mini-CRISPR array with a spacer targeting the *HHIV2_gp5* gene, including upstream and downstream flanking sequences for homologous recombination. **b**. Mini-CRISPR array spacers targeting the *HHIV2_gp5* gene. **c**. BREX defense activity against HHIV2 mutants with different HHIV2 *gp5* deletions. **VP1 protein sizes are calculated assuming Met103 as the translation start codon. **d**. Virion protein composition of wild-type and mutant HHIV2 viruses. Asterisks identify protein bands containing VP1 or its variants. **e**. A schematic overview of mutations identified in different HHIV2 mutant viruses within the upstream regulatory region and translation start site of the *HHIV2_gp5* gene. The ttc PAM of protospacer 0 is shown in red typeface.

To determine whether the VP1 protein is sufficient for BREX inhibition, we cloned SH1 *gp13* and HHIV2 *gp5*, along with 100 bp upstream sequences containing their promoter regions, generating plasmids SH1_P2_VP1 and HHIV2_P2_VP1. HHIV2_P2_VP1 inhibited BREX defence, while VP1 from SH1 did not (Fig. 2e, f). During the SH1 infection, *gp13* is continuously transcribed from three overlapping promoters along with downstream genes *gp14, gp15*, and *gp16* (31). A plasmid containing the entire *gp13-gp16* operon inhibited BREX^HAR^ and stop codons introduced into any one of the downstream genes resulted in the loss of BREX inhibition (Fig. 2f). These findings suggest that while VP1 from HHIV2 is sufficient for BREX repression, VP1 from SH1 requires involvement of Gp14, Gp15, and Gp16, which may explain its weaker inhibitory activity.

### Identifying HHIV2 VP1 protein regions required to overcome the BREX defense

We next attempted to knock out the *HHIV2_gp5* gene from the viral genome using the endogenous *H. hispanica* type I-B CRISPR-Cas system (32). The viral gene was targeted using a plasmid carrying a type I-B CRISPR mini-array, which included a *HHIV2_gp5*-specific spacer positioned between the two repeats. The array was transcribed from a constitutive *P*_*phar*_ promoter (33). The plasmid also carried *HHIV2_gp5* regions flanking the targeted site to provide a DNA repair template (Fig. 3a). The mini-array spacer targeted protospacer 1 inside the *HHIV2_gp5* gene (Fig. 3b). Screening of viral progeny produced upon infection of cells harbouring the mini-array plasmid showed that instead of expected viruses with completely deleted *HHIV2_gp5*, clones with partial deletions were recovered. Several individual HHIV2 mutants were isolated and sequenced. In every case, in-frame *HHIV2_gp5* deletions of various sizes were observed. Several individual viral isolates were next tested for their ability to overcome the BREX defense by titering on BREX+ and ΔBREX cells lawns (Fig. 3c, the mutants are identified by deleted VP1 amino acids using the revised numbering described above). A mutant with deletion spanning amino acids 61 to 879 (VP1Δ61-879) behaved as the wild-type virus and was insensitive to BREX. The VP1Δ38-966 mutant showed partial, ∼100-fold, protection from BREX; protection from VP1Δ21-844 was increased to ∼1000-fold. This level of BREX sensitivity is above that observed for sensitive viruses HHPV3 and HHPV4 and consistent with higher number of BREX^HAR^ recognition sites in the HHIV2 genome (Fig. 2a). Strikingly, VP1Δ23-892 efficiently infected BREX+ cells (10-fold protection). Overall, the results suggest that the VP1 segment containing amino acids 22-60 plays a critical role in overcoming the BREX defense.

We next targeted the remainder of the *HHIV2_gp5* gene of the VP1Δ21–844 virus using additional mini-array plasmids carrying spacers directed against protospacers 2 or 3 within *HHIV2_gp5* (Fig. 3b). In this way, we obtained double mutants, which again contained in-frame deletions. The VP1Δ21-844&Δ1052-1602 and VP1Δ21-844&Δ976-1364 double mutants were highly sensitive to BREX (100 000-fold less plaques on BREX+ compared to ΔBREX cell lawns, Fig. 3c). We therefore conclude that amino acids in the C-terminal part of VP1 also contribute to anti-BREX activity.

Since the complete deletion of the *HHIV2_gp5* gene would remove approximately one-sixth of the viral genome, our inability to completely delete this gene may have reflected the physical constraints on the amount of DNA that can be deleted without impairing the viral genome packaging (though the invariable appearance of in-frame deletions suggest that at least part of *HHIV2_gp5* is essential). To decrease *HHIV2_gp5* synthesis without grossly affecting the size of the viral genome, we used plasmids with spacer_-1 and spacer_0, targeting, respectively, the previously predicted or experimentally identified sequences at and around VP1 start codons (Fig. 3b). All escapers isolated on the lawn of cells expressing spacer_-1 contained mutations in the PAM sequence rather than in the protospacer, indicating that the targeted non-coding region ∼300 bp upstream of *HHIV2_gp5* reading frame is essential for productive infection. Escapers that emerged following the infection of *H. hispanica* cells expressing spacer_0, which targets the experimentally verified *HHIV2_gp5* start codon, carried either PAM mutations or small deletions within the protospacer sequence (Fig. 3e). One such mutant, designated “upstreamΔ1-2&VP1Δ1”, lost a sequence containing the start codon but acquired a 2 bp insertion in the beginning of the translated part of *HHIV2_gp5*. The resulting virus retained full ability to inhibit the BREX defense (Fig. 3e). We also identified another mutant, “upstreamΔ13-66”, with a 54-bp deletion. Compared to ΔBREX, the BREX+ cells were ∼10,000-fold resistant to this mutant (Fig. 3e). The final mutant, “upstreamΔ23-78”, harboured a similar upstream deletion but did not exhibit any BREX inhibition activity.

Virions of wild-type and several mutant viruses were purified and subjected to SDS PAGE analysis (Fig. 3d). A prominent high molecular weight VP1 band was observed in the wild-type virus sample. This band was less abundant in the upstreamΔ1-2&VP1Δ1 virion preparation. A smaller, ∼125 kDa band, containing VP1 peptides as determined by mass-spectrometric analysis was observed in VP1Δ38-966 virions preparation, consistent with the size of deletion. We were unable to identify the VP1 derivative in VP1Δ21-844&Δ1052-1602 virions by Coomassie staining. Likewise, virions with short upstream deletions (upstreamΔ13-66 and upstreamΔ23-78) had no detectable VP1 (Fig. 3d). The results demonstrate that VP1, a major protein in wild-type HHIV2 virions, is, surprisingly, not required for formation of infectious viral particles. The results also demonstrate that viruses containing VP1 with extensive deletions (VP1Δ38-966) and decreased abundance of VP1 (upstreamΔ1-2&VP1Δ1) can, respectively, partially or fully overcome the BREX defense.

## DISCUSSION

In this work, we demonstrate that the archaeal Type V BREX system of *H. hispanica* protects the host against a subset of haloviruses despite the fact that these viruses were originally isolated on a wild-type strain possessing this immune system. The apparent paradox can be explained by the emergence of rare viral variants that acquire host-specific BREX methylation during the initial rounds of infection, allowing them to subsequently replicate and overtake the population of initially protected cells. Consistent with this interpretation, propagation of the same viruses on cells lacking the BREX^HAR^ locus eliminated the protective modification, restoring the defense function of the *H. hispanica* BREX. Thus, as in bacterial systems, the outcome of viral infection depends not only on the viral genotype but also on the epigenetic state of the viral genome.

The recognition strategy of archaeal Type V system shown in our work resembles that of bacterial Type I BREX systems and differs from the mechanism proposed for the Pgl system (11–13). However, the *H. hispanica* BREX system has hallmarks that substantially expand the known diversity of BREX organisation and function. Most prominently, its locus encodes two active DNA methyltransferases, BrxX1 and BrxX2, that modify DNA independently. Methylation of sites recognized by both methyltransferases is required for full levels of protection against viral infection. Similar to Type I systems, deletion of either methyltransferase in the BREX^HAR^ system is not toxic for cells. The ability to generate such deletions can be explained by the involvement of methyltransferases in the restriction process: deletion of a methyltransferase gene abolishes not only the corresponding modification activity but also the associated restriction activity.

The archaeal system also differs from bacterial BREX by methylation chemistry and the composition of the modification module. In contrast to bacterial BREX systems, archaeal Type V system relies on methylation of cytosine residues in the host genome rather than adenines. In addition, all Type V loci encode two BrxC ATPases, and both copies are required for defense and DNA methylation, suggesting functional cooperation rather than redundancy. Finally, the last gene in the locus encodes an AAA+ ATPase, BrxHII, which shows no homology to *E. coli* BrxL. Similar to bacterial BREX systems, BrxHII appears to play a regulatory role but is not required for BREX-mediated DNA methylation.

Bacteriophages use diverse strategies to overcome bacterial BREX system. These include DNA-mimic proteins that bind the BREX methyltransferase BrxX, such as the Ocr protein from bacteriophage T7 (34), as well as proteins that disrupt BrxC multimerization and thereby inhibit BREX defense, demonstrated by the OrbA protein from the ICP1 phage (35). Another strategy is employed by T3 phage, which indirectly inhibits the BREX system through depletion of S-adenosylmethionine (SAM), a cofactor required for BREX methylation (36). In addition, bacteriophage T4 protects its genome by glycosylation of DNA, thereby allowing it to evade BREX-mediated immunity (3). Anti-BREX proteins are usually small, typically not exceeding 20 kDa. By contrast, here we demonstrate that some archaeal viruses can encode unusually large proteins for BREX inhibition: HHIV2 VP1 is 183 kDa and SH1 VP1 is 152 kDa. VP1 protein lacks homology to known proteins, except for VP1 from the closely related icosahedral virus PH1. In HHIV2, BREX inhibition relies solely on VP1, whereas in SH1 the smaller VP1 protein requires three additional viral proteins to achieve BREX inhibition.

We initially hypothesized that only a small domain of the VP1 protein might be responsible for anti-BREX activity, with the remaining regions required for viral replication. However, since multiple regions of VP1 can be deleted without abolishing viral viability, we propose that inhibition of the BREX system is the primary function of VP1. By targeting the upstream region of the *HHIV2_gp5* gene, we abolished VP1 synthesis and obtained an HHIV2 mutant lacking VP1. Our data suggest that the predicted start codon of VP1 is misannotated; nevertheless, this upstream region appears essential for the virus, as it could not be mutagenized without loss of viral viability.

The molecular mechanism of VP1-mediated BREX inhibition remains to be established. Notably, similar to the bacterial anti-BREX proteins Ocr (34) and OrbA (35), VP1 does not interfere with BREX-associated DNA methylation (Supplementary Fig.1), suggesting that VP1 selectively inhibits the defensive function of the BREX system rather than its modification activity.

In conclusion, these findings advance our understanding of prokaryotic immunity. We establish Type V BREX as an effective antiviral system in archaea, revealing a previously unrecognized dual-methyltransferase strategy for epigenetic discrimination and demonstrating that archaeal viruses encode specific anti-BREX proteins. Together, these results show that archaea and their viruses have evolved a molecular armory that is no less sophisticated than that described in bacteria, and highlight archaeal defense systems, along with their viral inhibitors, as a rich source of fundamentally new biology with significant potential for future biotechnological applications, as exemplified by CRISPR–Cas systems.

## DATA AVAILABILITY

The sequencing data have been deposited in the NCBI Sequence Read Archive (SRA) under BioProject accession number PRJNA1417484.

## ACKNOWLEDGMENTS

We would like to thank Prof. Hua Xiang (Institute of Microbiology, Chinese Academy of Science) for providing *Haloarcula hispanica* strains and HHPV2 virus. We thank Prof. Dennis Bamford for providing a collection of *H. hispanica* viruses. This research was funded by the Wellcome Trust (grant number 224665 to A. A. A.).

