## Supplementary materials for "A virion protein of an archaeal virus inhibits a Type V BREX defense system in *Haloarcula hispanica*"

### Supplementary Table S1. Strains and plasmids used in this work

| ***E. coli* strain** | **Comments** | **Source** |
|---|---|---|
| Stella | Strain for cloning | Lab stock |
| DH5α | Strain for cloning | Lab stock |
| ***Haloarcula hispanica* strain** | | |
| ATCC 33960 DF60 | Natural isolate with BREX$^{HAR}$ system with deleted *pyrF* | Hua Xiang |
| ATCC 33960 DF60 Δ*cas* | Natural isolate with BREX$^{HAR}$ system with deleted *pyrF* and *cas1-cas8* genes | Hua Xiang |
| ATCC 33960 DF60 ΔBREX | DF60 strain with deleted *pyrF* and BREX$^{HAR}$ system for CRISPR-targeting | This work |
| ΔBREX | DF60 Δ*cas* strain with deleted BREX$^{HAR}$ system | This work |
| ΔX1-HII | DF60 Δ*cas* strain with deleted *brxX1-Z-X2-HII* genes | This work |
| BREXΔN,<br>N = *brxA, B, C1, C2, X1, X2, Z, HII, nucS* | DF60 Δcas strain with a deleted BREX gene | This work |
| ***Haloarcula hispanica* viruses** | | |
| HHPV1 | | Dennis Bamford |
| HHPV2 | | Hua Xiang |
| HHPV3 | | Dennis Bamford |
| HHPV4 | | Dennis Bamford |
| SH1 | | Dennis Bamford |
| HHIV2 | | Dennis Bamford |
| HHTV1 | | Dennis Bamford |
| ***Haloarcula hispanica* plasmids** | | |
| pHAR | *pyrF*-based suicide vector for genome editing in *H. hispanica, pyrF, AmpR* | Hua Xiang |
| pHAR-ΔBREX | Plasmid for BREX$^{HAR}$ system deletion, *pyrF, AmpR* | This work |
| pHAR-ΔX1-HII | Plasmid for *X1-Z-X2-HI* genes deletion, *pyrF, AmpR* | This work |
| pHAR-ΔN,<br>N = *brxA, B, C1, C2, X1, X2, Z, HII, nucS* | Plasmid for deletion of individual *brx* genes, *pyrF, AmpR* | This work |
| pWL502 | *pyrF*-based vector of *H. hispanica, pyrF, AmpR* | Hua Xiang |
| pWL502-HHIV2_Pn, where n = 1 - 7 | Plasmids expressing HHIV-2 virus DNA fragments | This work |
| pWL502-HHIV2_P2_VP1 | Plasmids expressing *HHIV-2_gp5* gene only | This work |
| pWL502-HHIV2_P2_nStopVP1, n = 21, 90, 102, 107 | Plasmids expressing the HHIV-2_P2 fragment with a stop codon at different positions | This work |
| pWL502-SH1_P2 | Plasmid expressing SH1 virus *gp11-gp16* fragment | This work |
| pWL502-SH1_P2_gp13-16 | Plasmid expressing SH1 virus *gp13-gp16* fragment | This work |
| pWL502-SH1_P2_VP1 | Plasmid expressing *SH1_gp13* gene only | This work |
| pWL502-SH1_P2_StopVP1/GpN, N = 14, 15, 16 | Plasmids expressing the SH1_P2 fragment with a stop codon at different genes | This work |
| pWL502-HHIV2_VP1_spN, where N= -1, 0, 1, 2, 3, 4 | Plasmids expressing a spacer targeting different positions of *HHIV-2_gp5* gene | This work |

**Supplementary Table S2. Primers used in this work**

| | |
|---|---|
| US_ΔBREX _F | atataggatccgacatcgtgttgacgacgctgc |
| US_ΔBREX_R | gccgaagacgtgccctgctaaatcggtttctgaatgagactacctc |
| DS_ΔBREX_F | gaggtagtctcattcagaaaccgatttagcagggcacgtcttcggc |
| DS_ΔBREX_R | atataggtaccttttacgagtgcctggatcgacg |
| US_ΔX1-HII_F | atataggatcccacctgtaacccgaatcaggaga |
| US_ΔX1-HII_R | gaggtagtctcattcagaaaccgatacgagaatccgtgatgtttcaag |
| DS_ΔX1-HII_F | cttgaaacatcacggattctcgtatcggtttctgaatgagactacctc |
| DS_ΔX1-HII_R | atataggtaccttttacgagtgcctggatcgacg |
| US_ΔA_F | tatattctagaagtcctcgaagaaggcaacgt |
| US_ΔA_R | actcgacgcagccggtcgatgacacg |
| DS_ΔA_F | ccggctgcgtcgagtcgagcgacgg |
| DS_ΔA_R | tatatggtacctactcggagccgttgtcaacat |
| US_ΔB_F | tatattctagagagtatcgacggttacgaggg |
| US_ΔB_R | ttggagatgaagtcccggaagggacgt |
| DS_ΔB_F | ggacttcatctccaatgagcgacacac |
| DS_ΔB_R | tatatggtaccacacatcctcgaacgtctggc |
| US_ΔC1_F | tatatggatcctccctctattgtccagctccc |
| US_ΔC1_R | taataaccgtcggcagcgaaccagcc |
| DS_ΔC1_F | tgccgacggttattatccgatgacgacaaaacg |
| DS_ΔC1_R | tatatggtaccggccttcgatctggtgggc |
| US_ΔC2_F | tatatggatccggacttcacggatcggatcg |
| US_ΔC2_R | atcacagagaagatgtcgtgaatgtgtgtgtc |
| DS_ΔC2_F | catcttctctgtgatggtgatcgagtcgc |
| DS_ΔC2_R | tatatggtaccgttgcagaactcgtcaagcacc |
| US_ΔX1_F | tatatggatcctgacgcgcttgaatctcagg |
| US_ΔX1_R | cttccggcttgcgggggtgggtaga |
| DS_ΔX1_F | cccgcaagccggaagtcgttgaggacaag |
| DS_ΔX1_R | tatatggtacccttgttctagaagtttctctc |
| US_ΔX2_F | tatatggatccgaaccggcggaactcgct |
| US_ΔX2_R | cttccggatgagagtcagtgatattctcccag |
| DS_ΔX2_F | actctcatccggaagtggtggcggac |
| DS_ΔX2_R | tatatggtacccgacgaacaggacgcgg |
| US_ΔZ_F | tatatggatcctcaactcaaagagaccactatcatgacc |
| US_ΔZ_R | aaatcgagggcttcgggaagggtctgtttc |
| DS_ΔZ_F | cgaagccctcgatttcatcagcatcacgcag |
| DS_ΔZ_R | tatatggtacccaggctgtaggcggtcga |
| US_ΔHII_F | tatattctagaagacggtacaggagaacaaagacg |
| US_ΔHII_R | tagttctcttgcccggacgagaatgaagaatc |
| DS_ΔHII_F | cgggcaagagaactactgtttgactctcccatt |
| DS_ΔHII_R | tatatggtacccgtcgcgagacaagtcattcc |
| US_ΔnucS_F | tatatggatccgaggcctcgctctggtcg |
| US_ΔnucS_R | gagtatctccgactctaagcatgatgtcgaga |
| DS_ΔnucS_F | gagtcggagatactcggcgagactaccg |
| DS_ΔnucS_R | tatatggtaccgttcttccttgtcgagctggg |
| pWL502_lin_F | ggtaccgttttcttgtgtgcg |
| pWL502_lin_R | ggatccatgggggttgttgtc |
| HHIV2_P1_InF_F | accccccatggatcccactctctctctctctggggg |
| HHIV2_P1_InF_R | caagaaaacggtaccgccacttcattatcaactgtgggc |

| | |
|---|---|
| HHIV2_P2_InF_F | accccccatggatcctatcacggttttacaccgcgtga |
| HHIV2_P2_InF_R | caagaaaacggtacccgggtccgcgtcgatttg |
| HHIV2_P3_InF_F | accccccatggatccggccctatcacgcagaacgac |
| HHIV2_P3_InF_R | caagaaaacggtacccagcggactcgtcggagt |
| HHIV2_P5_InF_F | accccccatggatcctacaagcgtttggacgacga |
| HHIV2_P5_InF_R | caagaaaacggtacccatgacggaccaccctatcg |
| HHIV2_P6_InF_F | accccccatggatcccatataggccgctctgtggg |
| HHIV2_P6_InF_R | caagaaaacggtacccaacaccagcgacgaaatgg |
| HHIV2_P7_InF_F | accccccatggatccgtccgggtaaactccgggtt |
| HHIV2_P7_InF_R | caagaaaacggtaccactctctctctctctggggg |
| HHIV2_P2_21StopVP1_F | cgTGAaggTGAcgggtggaacc |
| HHIV2_P2_21StopVP1_R | cccgTCAcctTCAcgcccatgt |
| HHIV2_P2_90StopVP1_F | tacggTGacatggtcgattctaccc |
| HHIV2_P2_90StopVP1_R | gaccatgtCAccgtatcgttggtc |
| HHIV2_P2_102StopVP1_F | cccTgAatgagtgcgagcacag |
| HHIV2_P2_102StopVP1_R | cgcactcatTcAgggcactcc |
| HHIV2_P2_107StopVP1_F | cgagcTGagtgtacgacaccct |
| HHIV2_P2_107StopVP1_R | cgtacactCAgctcgcactcatg |
| HHIV2_P2_VP1_R | caagaaaacggtaccttcacacgtcctccgttgg |
| SH1_P2_InF_F | accccccatggatccacaagctccgagcggcgg |
| SH1_P2_InF_R | caagaaaacggtaccggtctgaccctatccaccct |
| SH1_P2_gp13_F | accccccatggatccaaacccggtgaatcccgg |
| SH1_P2_gp13_R | caagaaaacggtaccctacacctcctccgtcgg |
| SH1_P2_StopVP1_F | ccTGAaccTGAgagcacgacgac |
| SH1_P2_StopVP1_R | gctcTCAggtTCAggggtcgtcc |
| SH1_P2_StopGp14_F | TGAcaccgcgacTGAggcgacgag |
| SH1_P2_StopGp14_R | TCAgtcgcggtgTCAggatgggtcc |
| SH1_P2_StopGp15_F | TGAacggtcgaaTGActccccgaggg |
| SH1_P2_StopGp15_R | TCAttcgaccgtTCActcggtcacgg |
| SH1_P2_StopGp16_F | TGAgacgacctgTGAgccaccatcc |
| SH1_P2_StopGp16_R | TCAcaggtcgtcTCAgacctcgtcag |
| CRISPR_array_F1 | accccccatggatcccgaagggaacatatatgttactgcaggtacaacaccgagttaggaggtttcagacgaaccctcgt |
| CRISPR_array_R1 | acgagggttcgtctgaaacctcctaactcggtgttgtacctgcagtaacatatatgttcccttcgggatccatggggggt |
| CRISPR_array_F2 | cagacgaaccctcgtggggttgaagcgtttcagacgaagggtcgtgcccttgaagcaaaaaaaatctagaggtaccgtt |
| CRISPR_array_R2 | aacggtacctctagattttttttgcttcaagggcacgacccttcgtctgaaacgcttcaaccccacgagggttcgtctg |
| pWL502_crRNA_lin_F2 | ggttcccttgcgtacagttccc |
| pWL502_crRNA_lin_R2 | gcacacaagaaaacggtacctctagat |
| US_HHIV2_gp5_F | cgttttcttgtgtgcacgacgaccggaaccacg |
| US_HHIV2_gp5_R | tagaacccagttgccacttcattatcaactgtgg |
| DS_HHIV2_gp5_F | ggcaactgggttctacgacccaacggag |
| DS_HHIV2_gp5_R | gtacgcaagggaaccgcggcgacgacggtatagt |
| pWL502_crRNA_lin_F | gtttcagacgaagggtcgtgcc |
| pWL502_crRNA_lin_R | ctcctaactcggtgttgtacctgc |
| spacer_-1_F | acaccgagttaggaggtttcagacgaaccctcgtggggttgaagctatgcccacagttgataatgaagtggcaacttcgac gtttcagacgaaggg |
| spacer_-1_R | cccttcgtctgaaacgtcgaagttgccacttcattatcaactgtgggcatagcttcaaccccacgagggttcgtctgaaacctcc taactcggtgt |
| spacer_0_F | acaccgagttaggaggtttcagacgaaccctcgtggggttgaagctaccctgcgccgggagtgcccggcatgagtgcgagc gtttcagacgaaggg |
| spacer_0_R | cccttcgtctgaaacgctcgcactcatgccgggcactcccggcgcagggtagcttcaaccccacgagggttcgtctgaaacc tcctaactcggtgt |
| spacer_1_F | acaccgagttaggaggtttcagacgaaccctcgtggggttgaagccacgaggtcggcggccagtatcaacaggacgcccta gtttcagacgaaggg |

| | |
|---|---|
| spacer_1_R | cccttcgtctgaaactagggcgtcctgttgatactggccgccgacctcgtggcttcaaccccacgagggttcgtctgaaacct cctaactcggtgt |
| spacer_2_F | acaccgagttaggaggtttcagacgaaccctcgtggggttgaagctaaacaggtacggcagcctcgaacgggttgcgacca gtttcagacgaaggg |
| spacer_2_R | cccttcgtctgaaactggtcgcaacccgttcgaggctgccgtacctgtttagcttcaaccccacgagggttcgtctgaaacctc ctaactcggtgt |
| spacer_3_F | acaccgagttaggaggtttcagacgaaccctcgtggggttgaagcgagccgaccggcggcgaccaaggcacgccggctacg gtttcagacgaaggg |
| spacer_3_R | cccttcgtctgaaaccgtagccggcgtgccttggtcgccgccggtcggctcgcttcaaccccacgagggttcgtctgaaacct cctaactcggtgt |
| spacer_4_F | acaccgagttaggaggtttcagacgaaccctcgtggggttgaagcgacatggccgcgaggtcggcgtcgagcagggtatc gtttcagacgaaggg |
| spacer_4_R | cccttcgtctgaaacgataccctgctcgacgccgacctcgcggcccatgtcgcttcaaccccacgagggttcgtctgaaacct cctaactcggtgt |

**Supplementary Figure 1.** Defense activity against HHPV3 (right column) and HHPV4 (left column) viruses propagated on BREX+ HHIV2_P2 lawn.

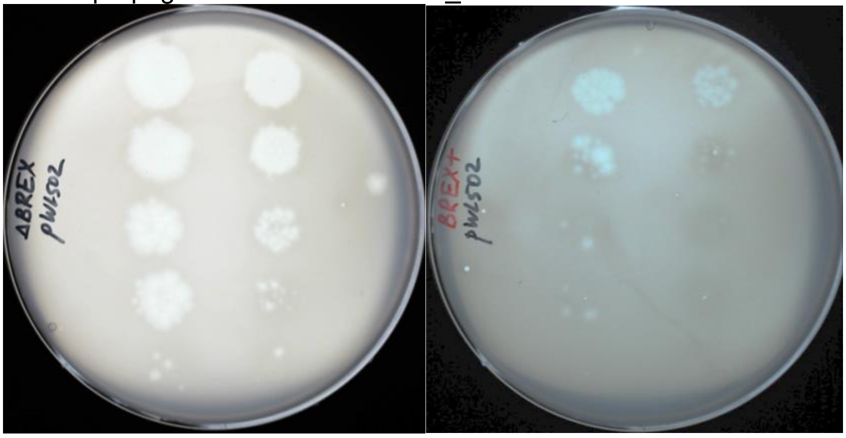
